# A two-year study on the phenology, host and habitat associations, and pathogens of *Haemaphysalis longicornis* in Virginia, U.S.A.

**DOI:** 10.1101/2021.01.20.427350

**Authors:** Alec T. Thompson, Seth A. White, David Shaw, Kayla B. Garrett, Seth T. Wyckoff, Emily E. Doub, Mark G. Ruder, Michael J. Yabsley

**Affiliations:** Southeastern Cooperative Wildlife Disease Study, Department of Population Health, College of Veterinary Medicine, University of Georgia, Athens GA; Center for the Ecology of Infectious Diseases, Odom School of Ecology, University of Georgia, Athens GA; Warnell School of Forestry and Natural Resources, University of Georgia, Athens GA

**Author notes:** Corresponding authors: Alec T. Thompson, 589 D.W. Brooks Dr., Athens GA 30602,; and Michael J. Yabsley, 589 D.W. Brooks Dr., Athens GA 30602.

**Keywords:** *Haemaphysalis longicornis*, phenology, seasonal abundance, host associations, habitat associations

## Abstract

Understanding the abiotic and biotic variables affecting tick populations is essential for studying the biology and health risks associated with vector species. We conducted a study on the phenology of exotic *Haemaphysalis longicornis* (Asian longhorned tick) at a site in Albemarle County, Virginia. We assessed the importance of available wildlife hosts, habitats, and microclimate variables such as temperature, relative humidity, and wind speed on this exotic tick’s presence and abundance. In addition, we determined the prevalence of selected tick-borne pathogens potentially transmitted by *H. longicornis*. We determined that the seasonal activity of *H. longicornis* was slightly different from previous studies in the northeastern United States. We observed nymphal ticks persist year-round but were most active in the spring, followed by a peak in adult activity in the summer and larval activity in the fall seasons. We also observed a lower probability of detecting *H. longicornis* in field habitats and the summer months. In addition, we detected *H. longicornis* on several wildlife hosts, including coyote (*Canis latrans*), eastern cottontail (*Sylvilagus floridanus*), raccoon (*Procyon lotor*), Virginia opossum (*Didelphis virginiana*), white-tailed deer *(Odocoileus virginianus*), woodchuck (*Marmota monax*), and a *Peromyscus* sp. This is the first detection of this tick on a rodent host important to the epidemiology of tick-borne pathogens of humans and animals. Finally, we continued to detect the exotic piroplasm parasite, *Theileria orientalis* Ikeda, in *H. longicornis* as well as other pathogens, including *Rickettsia felis, Anaplasma phagocytophilum* (AP-1), and a *Hepatozoon* sp. previously characterized in *Amblyomma americanum*. These represent some of the first detections of arthropod-borne pathogens native to the United States in host-seeking *H. longicornis*. These data increase our understanding of *H. longicornis* biology in the United States and provide valuable information into the future health risks associated with this tick and pathogens.

## Introduction

The seasonal abundance of tick populations has implications for pathogen transmission to humans and animals. Abiotic factors, habitat conditions, and the presence of host species acting as tick hosts and pathogen reservoirs all strongly influence the risk of transmission (Gleim et al., 2014; MacDonald, 2018). In addition, understanding habitat preferences of ticks coupled with presence/absence data of different species can provide critical information to identify high-risk regions for certain tick-borne diseases. Therefore, data on the peak activity periods and habitat preferences of tick species are needed to better understand the seasonal risk for pathogen transmission and the deployment of tick mitigation strategies.

In the United States, the biology for most native tick species relevant to human and companion animal health are well studied. However, the recent widespread detections of exotic *Haemaphysalis longicornis* (Asian longhorned tick) in the United States raise concern for native and exotic tick-borne pathogen transmission, warranting further investigation (Beard et al., 2018; Hutcheson et al., 2018). Native to East Asia, *H. longicornis* has been introduced into multiple regions of the world, including Australia, New Zealand, and now the United States, where it is a recognized One Health concern (Beard et al., 2018; Heath, 2020; Hoogstraal et al., 1968). Although *H. longicornis* was first detected outside of quarantine zones in New Jersey in late 2017, an examination of archived ticks indicated this species has been in the United States since as early as 2010 in West Virginia (Beard et al., 2018; Rainey et al., 2018). Recent molecular analysis indicates that at least three females of different lineages were introduced (Egizi et al., 2020). To date, *H. longicornis* has been detected in multiple states in the eastern United States and has been found on many different hosts species raising concerns for potential pathogen transmission between humans, domestic animals, and wildlife species (Beard et al., 2018; Duncan et al., 2020; Tufts et al., 2019; USDA-APHIS-VS, 2021; White et al., 2020).

Many studies regarding the biology and seasonal abundance of *H. longicornis* have been conducted within its established range outside of the United States (Heath, 2016; Zheng et al., 2012). Generally, *H. longicornis* life stages follow a seasonal trend where adults are active in the summer, followed by a peak in larvae in the fall with nymphal ticks overwintering and becoming active again in the spring. Currently, few phenology and habitat association studies for *H. longicornis* have been completed in the United States (Bickerton et al., 2020; Piedmonte et al., 2020; Tufts et al., 2020a, 2019). While the life stage trends follow what has previously been reported outside of the United States, these studies were short term or situated in suburban areas and were limited to the northern regions of the recognized *H. longicornis* range in the United States. Thus, additional work on long-term seasonal trends in areas with a greater diversity of hosts is needed to better understand the phenology and habitat and host associations of *H. longicornis* in the United States.

In addition to phenological studies, there are few data on the abundance and diversity of pathogens in *H. longicornis* in the United States. Most recently, a study from Pennsylvania has detected *Borrelia burgdorferi* sensu stricto (causative agent for Lyme Disease) in a single *H. longicornis* (Price et al., 2021). Another study tested host-seeking ticks in Virginia and detected an exotic cattle pathogen, *Theileria orientalis* Ikeda strain (Thompson et al., 2020b). Finally, a study from New York using a shotgun sequencing approach failed to detect any bacterial or viral pathogens from *H. longicornis* (Tufts et al., 2020b). However, within its established range outside of the United States, *H. longicornis* is associated with numerous pathogens of both human and veterinary concern including *Anaplasma*, *Theileria*, *Babesia*, and spotted fever group *Rickettsia* spp. (Hong et al., 2019; Kang et al., 2016; Lee et al., 2003; Lee and Chae, 2010; Lu et al., 2013; Sivakumar et al., 2014; Watts et al., 2016). Although the role that this tick will play in the transmission of native tick-borne pathogens is currently unclear, the introduction of *H. longicornis* to the United States is of great concern to human and veterinary health as it could potentially alter the dynamics of endemic diseases and enable the transmission and potential establishment of exotic pathogens.

This study aims to obtain data on the phenology, habitat, and host associations for *H. longicornis* and the prevalence of selected pathogens in host-seeking *H. longicornis* relevant to both public and veterinary perspectives. To accomplish this, a multi-year study at a single site in rural Virginia was conducted where ticks were systematically collected from the environment and wildlife hosts.

## Methods

### General

Wildlife and ticks were sampled seasonally at a 109-acre cattle operation with a known *H. longicornis* infestation in Albemarle County, Virginia, from May 2019 to September 2020. This site also previously experienced cattle mortalities caused by the exotic intraerythrocytic parasite *Theileria orientalis* Ikeda genotype (Oakes et al., 2019). Approximately 72% of the property is a mixture of cow-calf beef cattle grazing and hay production pastures, 25% is a small plot of hardwood forest with a high mixed brush understory. The remaining 3% consists of owner residences. The surrounding properties are of similar habitats used for pastures or crops. To investigate seasonal phenology for *H. longicornis*, host and habitat associations, we conducted two 5-day sampling periods during May (spring) and September (fall) (rodent trapping and environmental sampling), and a 12-day sampling period during July for the summer season (rodent, meso-mammal, and environmental sampling). All collected ticks are stored in 70% ethanol for morphologic identification using published keys (Clifford et al., 1961; Cooley and Kohls, 1944; Durden and Keirans, 1996; Egizi et al., 2019; Keirans and Litwak, 1989; Walker, 2000). Suspect *H. longicornis* were confirmed using molecular techniques as described (Thompson et al., 2020a).

### Wildlife sampling

Rodents and meso-mammals were trapped using methods as described (White et al., 2020). Briefly, Sherman box traps (H. B. Sherman Inc., Tallahassee, FL) baited with peanut butter cereal were used for rodents, and Havahart cage traps (Woodstream Corporation, Lititz, PA) baited with either canned dog food or sardines were used for meso-mammals such as Virginia opossums (*Didelphis virginiana*) and raccoons (*Procyon lotor*). Traps were preferentially placed in areas to avoid tampering by cattle and to maximize capture success. Rodents were trapped during all sampling periods, but meso-mammals were only trapped during the summer sampling period. Other wildlife species, such as white-tailed deer (*Odocoileus virginianus)*, coyote (*Canis latrans*), reptiles, and amphibians were opportunistically sampled when available (e.g., vehicle collision or manual capture). All meso-mammals except for small rodent species and Virginia opossums were immobilized prior to tick collection using the premixed combination of nalbuphine (40 mg/ml), azaperone (10 mg/ml), and medetomidine (10 mg/ml) (NalMed; ZooPharm, Laramie, Wyoming USA) administered by intramuscular (IM) injection (0.3 mg/kg). Anesthetized animals were reversed with 0.6 mg atipamezole/kg and 0.15 mg naltrexone/kg (ZooPharm) IM and released at the site of capture after recovery. All animal capture and handling techniques were reviewed and approved by the University of Georgia’s Institutional Animal Care and Use Committee (A2018 06-027).

### Environmental sampling

Tick drags were conducted during the same sampling periods as the wildlife trapping. Host-seeking ticks were collected from field, forest, and edge habitats via tick drags using a 1m^2^ felt cloth. Field habitats primarily consisted of switchgrass (*Panicum virgatum*) and pastures used for cattle grazing or hay production. Forest habitats consisted of hardwood forest with a high understory, with groundcover primarily of invasive Japanese stiltgrass (*Microstegium vimineum*) and wild blackberry (*Rubus fruticosus*). Edge habitats contained previously listed grasses and beefsteak mint (*Perilla frustescens*) and followed pasture and forest fence line. Flagging on the fence line’s forest side was done when possible. Each drag was 100m long, with stops every 10-20 m for tick removal. During the sampling period, each habitat was sampled six times daily except the forest habitat (sampled four times) due to limited space in this habitat. Tick dragging was generally conducted in the early afternoon or later in the evening if the ground was still wet. Dragging was not conducted on rainy days. Microclimate data such as average wind speed, temperature, and relative humidity were collected at the beginning of every drag using a Kestrel 3000 wind meter (Nielsen-Kellerman Company, Boothwyn, PA).

### Analysis of Haemaphysalis longicornis phenology and habitat associations

Each life stage’s seasonal pattern of activity was visualized in R (version 3.6.2 – R Core Team 2018). For questing tick phenology data, counts of *H. longicornis* from all drags were pooled for a given collection day, and the proportion of each life stage (larvae, nymphs, and adults) was calculated and plotted. A best fit curve was generated between sampling periods using generalized additive models (GAM) fitted to the count data across time (Moussus et al., 2009). General linearized models (GLMs) were used to determine which abiotic variables were the most important predictors for *H. longicornis* abundance and presence. The variables tested were habitat type (field, forest, and edge), season (spring, summer, fall), and collected microclimate variables (average wind speed, temperature, and relative humidity). For the tick presence models, a logit link function with a binomial response variable was used, whereas the tick abundance models used a Poisson regression.

### Pathogen surveillance

Only adult and nymphal ticks collected from the environment were screened for pathogens. Ticks were medially bisected with a sterile razor blade and then dried overnight in a sterile hood to allow ethanol to evaporate. DNA was extracted from one half using the Qiagen DNeasy Blood and Tissue Kit (Qiagen, Germantown, MD) following the manufacturer’s protocol. The other half of the tick was preserved in 70% ethanol for archiving and morphologic identification. Both native ticks and exotic *H. longicornis* were screened for selected pathogens, including *Theileria, Babesia*, *Hepatozoon, Ehrlichia*, *Anaplasma*, *Rickettsia*, and *Borrelia* spp. using published polymerase chain reaction (PCR) protocols (Table 1). Positive PCR amplicons were visualized on 2% agarose gels stained with GelRed (Biotium, Hayward, CA, USA). Amplicons were purified from gels using the QIAquick gel extraction kit (Qiagen, Hilden, Germany) and submitted for bi-directional Sanger sequencing at the Genewiz Corporation (South Plainfield, NJ, USA). Chromatograms were analyzed using Geneious R11 (Geneious, Auckland, New Zealand, https://www.geneious.com). For the piroplasm surveillance, the genotype of *Theileria orientalis* detected was determined by sequence analysis of the major piroplasm surface protein (*MPSP*) primer gene. For *Anaplasma phagocytophilum* screening, all ticks were initially screened with the major surface protein-2 (*MSP-2*) primer pair, and a 16S rRNA primer set was used to type variants (human (AP-ha) or white-tailed deer (AP-1)) detected. All unique sequences obtained from this study were deposited to GenBank under the Accession numbers: MW480558, MW491252-MW491253.

**Table 1.**
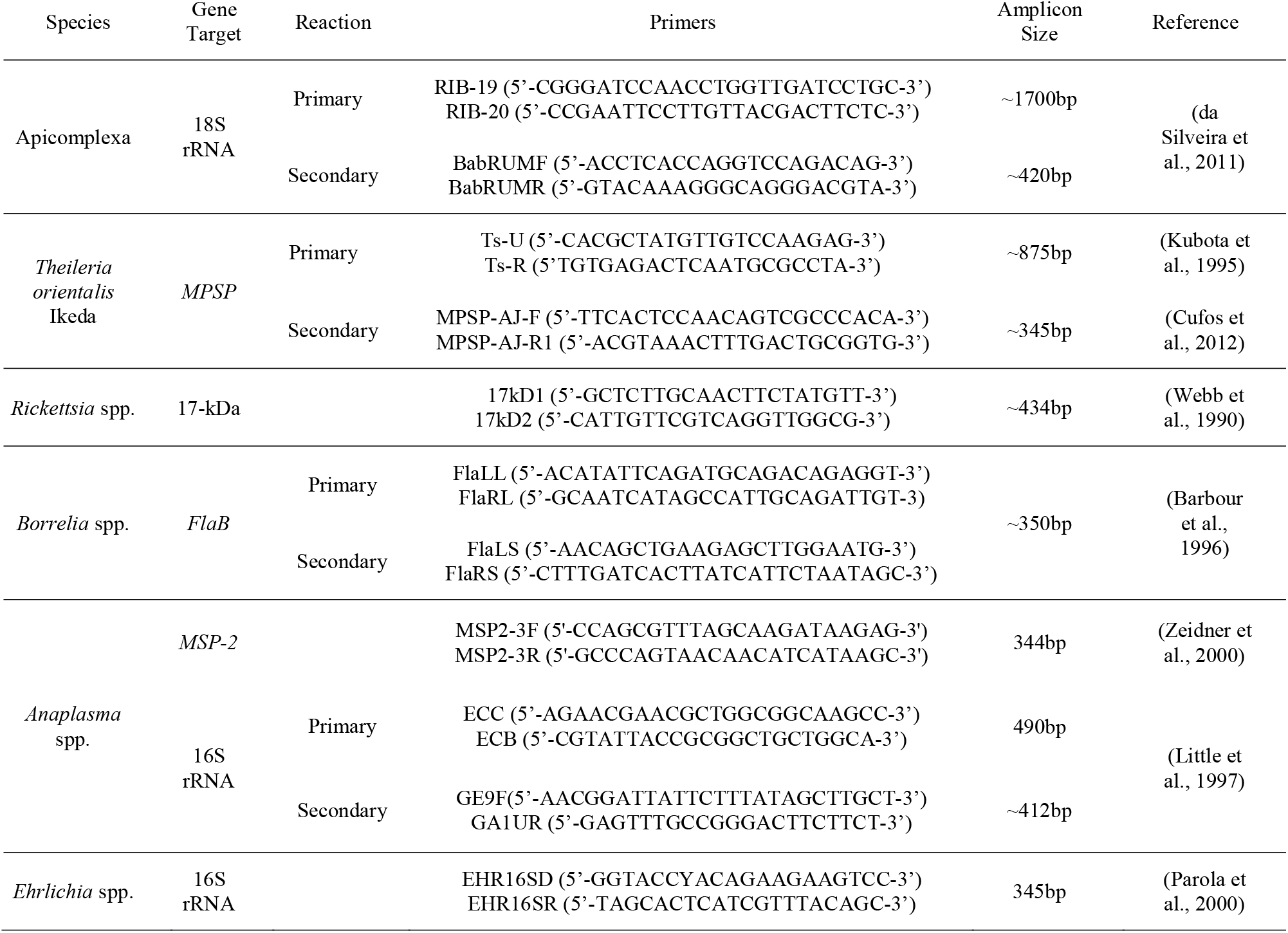
Gene targets and PCR primers used to detect pathogens in host-seeking ticks.

## Results

Over the two sampling years, 1582 ticks were collected from 203 hosts and 478 drags (Table 2, Figure 1). From the wildlife sampling, a total of 670 ticks were collected, with the most abundant tick species being *Dermacentor variabilis* (n=205), followed by *Amblyomma americanum* (n=152) and *Amblyomma maculatum* (n=133). Most *Ixodes* species collected from the wildlife hosts were *Ixodes scapularis* (n=92), but lower numbers of *Ixodes cookei* (n=31) and *Ixodes texanus* (n=10) were collected. The native rabbit tick, *Haemaphysalis leporispalustris* (n=4), was the least abundant tick collected from wildlife (Table 2, Figure 1). A total of 43 *H. longicornis* were collected from 18 different individual hosts of the following species: coyote, eastern cottontail (*Sylvilagus floridanus*), raccoon, Virginia opossum, white-tailed deer, woodchuck (*Marmota monax*), and a *Peromyscus* sp. (Table 2, Figure 1). Two *H. longicornis* nymphs were also opportunistically collected from humans as part of daily tick checks.

**Table 2.** Summary of ticks collected from hosts and drags from May 2019 to September 2020.^a^

| Sampling | Species | n | # infested with ticks | # with ALT<sup>b</sup> | Tick species identified |
| --- | --- | --- | --- | --- | --- |
| Spring 2019<br>May 2 – May 8 | Gray squirrel | 1 | 0 | 0 | - |
|  | Hispid cotton rat | 1 | 0 | 0 | - |
|  | *Peromyscus* sp. | 41 | 22 (54%) | 0 | *A. americanum*, *A. maculatum*, *I. scapularis* |
|  | Garter snake | 1 | 0 | 0 | - |
|  | Tick drag | 64 | 31 (48%) | 29 (45%) | *A. americanum*, *D. variabilis*, *H. leporispalustris*, *H. longicornis* |
| Summer 2019<br>July 9 – July 21 | Alleghany woodrat | 1 | 0 | 0 | - |
|  | Eastern cottontail | 1 | 1 (100%) | 0 | *H. leporispalustris* |
|  | Gray squirrel | 2 | 1 (50%) | 0 | *D. variabilis* |
|  | *Peromyscus* sp. | 17 | 1 (6%) | 0 | *I. scapularis* |
|  | Raccoon | 9 | 9 (100%) | 4 (44%) | *A. americanum*, *A. maculatum*, *D. variabilis*, *H. longicornis*, *I. cookei*, *I. scapularis*, *I. texanus* |
|  | Virginia opossum | 6 | 6 (100%) | 1 (17%) | *A. americanum*, *D. variabilis*, *H. longicornis*, *I. scapularis* |
|  | White-tailed deer | 1 | 1 (100%) | 1 (100%) | *A. americanum*, *H. longicornis*, *I. scapularis* |
|  | Woodchuck | 2 | 1 (50%) | 1 (50%) | *H. longicornis* |
|  | Human | 2 | 1 (50%) | 0 | *A. americanum*, *D. variabilis* |
|  | Tick drag | 110 | 25 (23%) | 18 (16%) | *A. americanum*, *D. variabilis*, *H. longicornis* |
| Fall 2019<br>Sept. 5 – Sept. 11 | Alleghany woodrat | 1 | 0 | 0 | - |
|  | Coyote | 2 | 2 (100%) | 2 (100%) | *D. variabilis*, *I. scapularis*, *H. longicornis* |
|  | *Peromyscus* sp. | 9 | 1 (11%) | 0 | *A. americanum* |
|  | Human | 2 | 2 (100%) | 1 (50%) | *A. americanum*, *H. longicornis* |
|  | Tick drag | 64 | 33 (52%) | 30 (47%) | *A. americanum*, *H. leporispalustris*, *H. longicornis* |
| Spring 2020<br>May 3 – May 9 | Hispid cotton rat | 1 | 1 (100%) | 0 | *A. americanum* |
|  | *Peromyscus* sp. | 30 | 22 (73%) | 0 | *A. maculatum*, *D. variabilis*, *I. scapularis* |
|  | Woodland jumping mouse | 1 | 1 (100%) | 0 | *A. americanum* |
|  | Human | 1 | 1 (50%) | 0 | *A. americanum*, *D. variabilis* |
|  | Tick drag | 64 | 25 (39%) | 21 (33%) | *A. americanum*, *D. variabilis*, *H. longicornis*, *I. scapularis* |
| Summer 2020<br>July 6 – July 19 | Alleghany woodrat | 3 | 0 | 0 | - |
|  | Eastern cottontail | 1 | 1 | 1 (100%) | *H. leporispalustris*, *H. longicornis* |
|  | Hispid cotton rat | 1 | 0 | 0 | - |
|  | *Peromyscus* sp. | 23 | 7 (30%) | 1 (4%) | *A. maculatum*, *H. longicornis*, *I. scapularis* |
|  | Raccoon | 4 | 4 (100%) | 1 (25%) | *A. americanum*, *D. variabilis*, *H. longicornis*, *I. cookei*, *I. scapularis*, *I. texanus* |
|  | Virginia opossum | 10 | 8 (80%) | 4 (44%) | *A. americanum*, *D. variabilis*, *H. leporispalustris*, *H. longicornis*, *I. scapularis* |
|  | Woodchuck | 2 | 1 (50%) | 0 | *A. americanum* |
|  | House sparrow | 1 | 0 | 0 | - |
|  | American toad | 1 | 0 | 0 | - |
|  | Broadheaded skink | 1 | 0 | 0 | - |
|  | Eastern ratsnake | 2 | 0 | 0 | - |
|  | Human | 2 | 2 (100%) | 1 (50%) | *A. americanum*, *D. variabilis*, *H. longicornis*, *I. scapularis* |
|  | Tick drag | 128 | 52 (41%) | 17 (13%) | *A. americanum*, *D. variabilis*, *H. longicornis*, *I. scapularis* |
| Fall 2020<br>Sept. 6 – Sept. 12 | Hispid cotton rat | 7 | 4 (57%) | 0 | *A. maculatum*, *I. scapularis* |
|  | *Peromyscus* sp. | 8 | 2 (25%) | 0 | *A. maculatum*, *I. scapularis* |
|  | Virginia opossum | 2 | 2 (100%) | 0 | *I. scapularis* |
|  | Carolina wren | 1 | 0 | 0 | - |
|  | Garter snake | 1 | 0 | 0 | - |
|  | S. Leopard frog | 1 | 0 | 0 | - |
|  | Tick drag | 48 | 20 (42%) | 20 (42%) | *A. americanum*, *H. longicornis* |

<sup>a</sup> Raw data describing total number of species and life stages for each host and drag sampled can be found in the supplementary data.
<sup>b</sup> ALT, Asian longhorned tick

**Figure 1.**
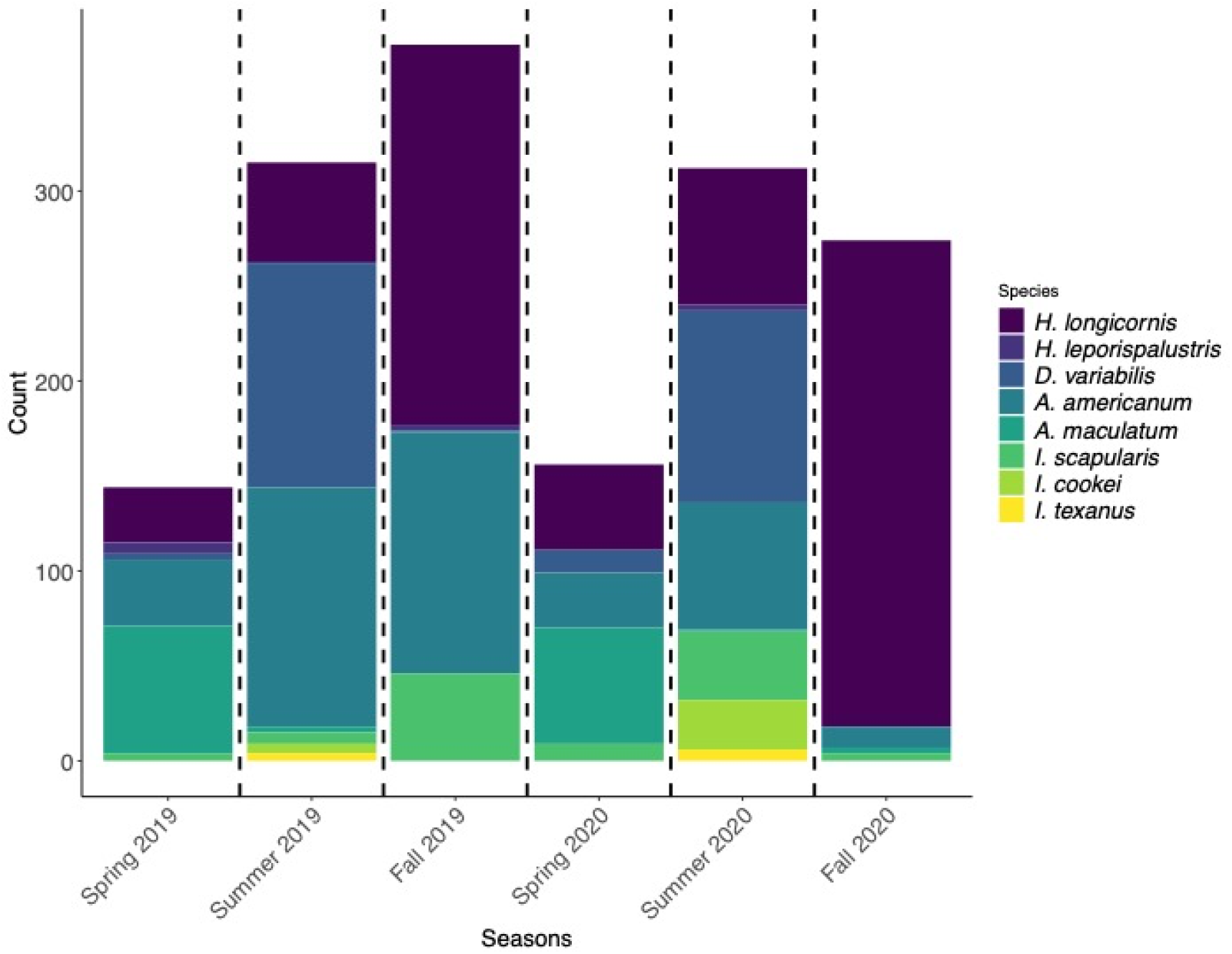
Diversity and abundance of ticks collected during the sampling periods from 2019-2020.

A total of 912 ticks were collected during environmental sampling. Of those, a majority were *H. longicornis* (n=615) followed by *A. americanum* (n=248), *D. variabilis* (n=30), *I. scapularis* (n=14), and *H. leporispalustris* (n=5) (Table 2, Figure 1). *Haemaphysalis longicornis* was collected from every habitat type sampled (field, forest, and edge). There was no significant difference in forest and edge habitats, but we observed a lower probability to find *H. longicornis* in field habitats (p < 0.05, Figure 2A). Season also had a significant effect on the probability of occurrence as we were less likely to detect *H. longicornis* in the summer season (p < 0.001, Figure 2B). There was no significant effect of the abiotic variables (average wind speed, temperature, and relative humidity) measured on *H. longicornis* presence or abundance. For phenology, nymphs were found in every season but were most active in the spring. This spring peak in nymphs is followed by a smaller adult peak in summer, followed by a large larval peak in the fall (Figure 3).

**Figure 2.**
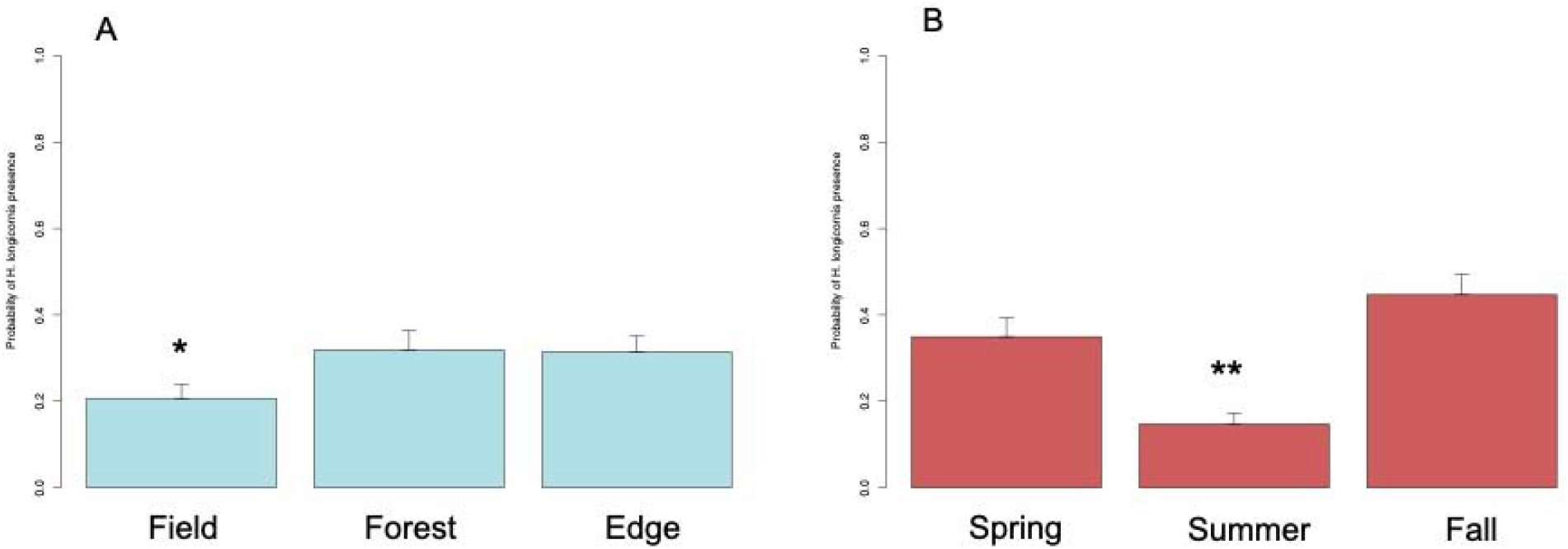
Effects of environmental variables on *Haemphysalis longicornis* presence. A) There is a significant effect of habitat type on *H. longicornis* presence. It is less likely to find *H. longicornis* in field habitats (*, p < 0.05) when compared to forest or edge habitats. B) There is a significant effect of season on *H. longicornis* presence. It is less likely to find *H. longicornis* during the summer (**, p < 0.001) when compared to spring or fall.

**Figure 3.**
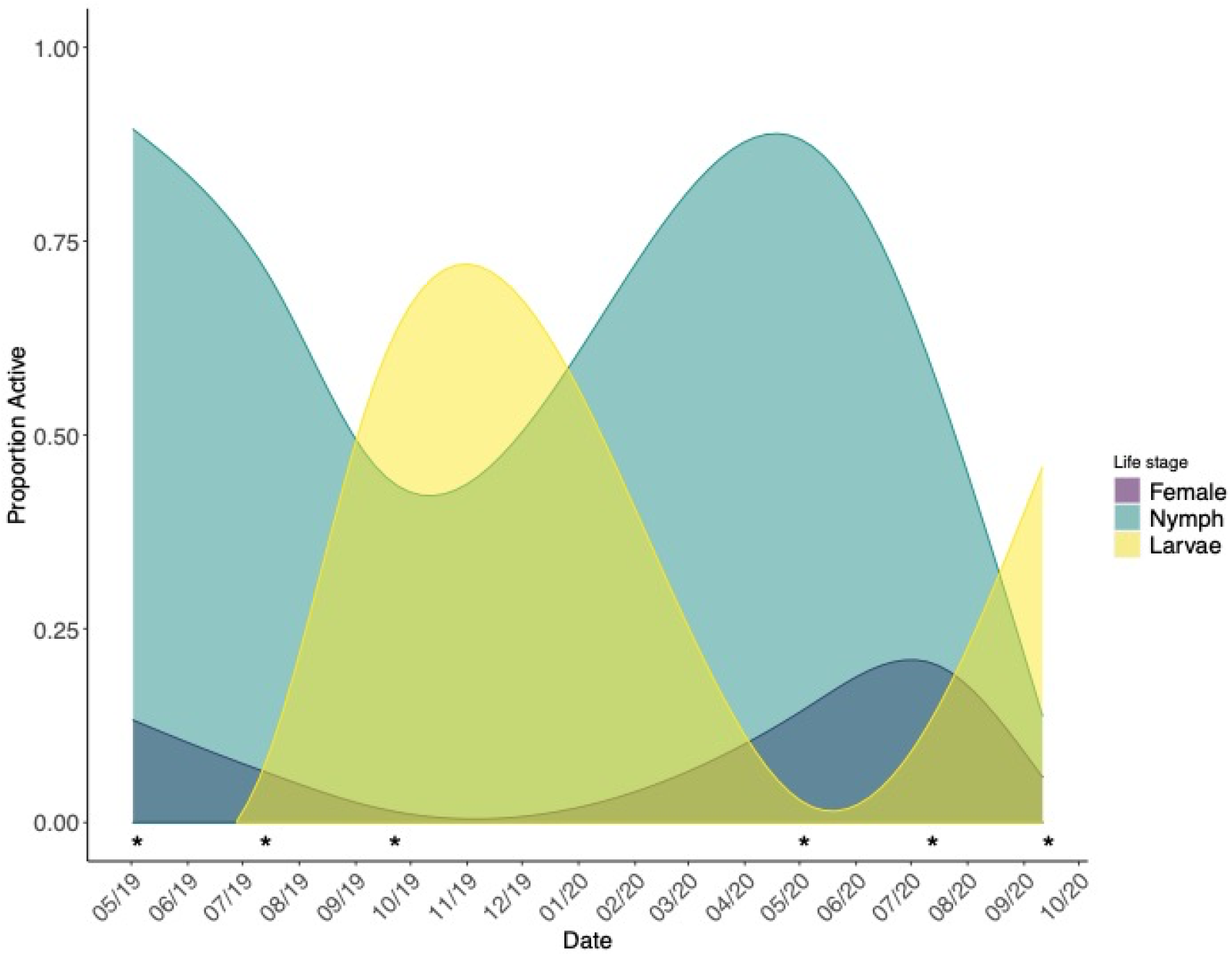
Seasonal activity of different *H. longicornis* life stages from 2019-2020. Nymphs (blue) are most active during the spring, with adults becoming active in summer (purple) and larvae becoming active in the fall (yellow). Asterisks indicate start of seasonal sampling periods. The adult peak was missed during the 2019 sampling period.

All host-seeking nymphal and adult ticks collected during this study (n=410) were screened for selected pathogens relevant to human and veterinary health (Table 1). However, our primary interest was exotic *Theileria orientalis* Ikeda genotype and native pathogens present in host-seeking *H. longicornis. Theileria orientalis* was detected in *H. longicornis* during both the 2019 and 2020 sampling periods (Table 3; Thompson et al., 2020b). Sequence analysis of partial *MPSP* gene sequences of all *T. orientalis* samples were 100% identical to the Ikeda genotype (JQ781070). In 2019, a single *H. longicornis* nymph was positive for *Rickettsia felis* (100% to MK509751), and two nymphs from 2020 were positive for a *Hepatozoon* sp. (100% to MT259335) (Table 3). Several *H. longicornis* nymphs from 2019 (n=1) and 2020 (n=7) were positive for *A. phagocytophilum* (99.4-100% to CP006617) (Table 3). Two of these *A. phagocytophilum* positive ticks were also positive with the 16S rRNA gene PCR (100% to MK341075), and the two nucleotide polymorphisms at bases 76 and 84 were consistent with *A. phagocytophilum* variant 1 (AP-1) associated with white-tailed deer (Dugan et al., 2006; Massung et al., 2003).

**Table 3.** Results of pathogen screening from host-seeking ticks collected from Albemarle Co., Virginia

| Species | Life stage<sup>a</sup> | Apicomplexa | Apicomplexa Species<sup>b</sup> | *Anaplasma phagocytophilum*<sup>c</sup> | *Borrelia* spp. | *Ehrlichia* spp. | *Rickettsia* spp.<sup>e</sup> |
| --- | --- | --- | --- | --- | --- | --- | --- |
| *Amblyomma americanum* | 13/12/96 | 2/2/13 | *Theileria cervi* (n=3),<br>*Babesia* spp. Coco (n=1),<br>*Hepatozoon* sp. (n=13) | 0/1/4 | 0/1/0 | 3/1/4<sup>d</sup> | 2/1/16 |
| *Dermacentor variabilis* | 15/15/0 | 0/0/0 | - | 0/0/0 | 0/0/0 | 0/0/0 | 1/1/1 |
| *Haemaphysalis leporispalustris* | 0/1/1 | 0/0/0 | - | 0/0/0 | 0/0/0 | 0/0/0 | 0/0/0 |
| *Haemaphysalis longicornis* | 0/18/229 | 0/0/24 | *Theileria orientalis* (n=22), *Hepatozoon* sp. (n=2) | 0/0/8 | 0/0/0 | 0/0/0 | 0/2/3 |
| *Ixodes scapularis* | 0/0/10 | 0/0/2 | *Babesia odocoilei* (n=2) | 0/0/0 | 0/0/1 | 0/0/0 | 0/0/8 |
| Total (410) | 28/46/336 | 2/2/39 |  | 0/1/12 | 0/1/1 | 3/1/4 | 3/4/28 |

<sup>a</sup> Life stages and results are represented as the number of Male/Female/Nymph tested or positive, respectively.
<sup>b</sup> All *Theileria orientalis* were confirmed as the Ikeda genotype using a supplemental PCR targeting the *MPSP* gene (see Table 1), some results are previously described in Thompson et al. 2020b.
<sup>c</sup> Only a single nymph of the *A. americanum* and two nymphs of *H. longicornis* were PCR positive for the 16S rRNA gene target, these were determined to the AP-1 variant (non-human pathogen).
<sup>d</sup> A single adult male and female were both infected with *E. ewingii* (100% to U96436) and a nymphal stage was infected with *E. chaffeensis* (100% to NR074500), others were infected with ‘*Candidatus* Midichloria mitochondrii’.
<sup>e</sup> All *Rickettsia* spp. detected were endosymbionts of ticks, except for a single *Rickettsia felis* (100% to MK509751) was detected in a *H. longicornis* nymph.

No *T. orientalis* Ikeda was detected in any native tick species screened, but numerous native piroplasm species relevant to veterinary health were detected from the 2019 and 2020 sampling periods. Data for the piroplasm screening from 2019 has been previously reported (Thompson et al., 2020b), but briefly, we detected a *Theileria* sp. of white-tailed deer (often called *T. cervi*) (n=3; 99% to AY35135), *Babesia* spp. Coco (n=1; 99% to EU109716), and a *Hepatozoon* sp. (n=1; 99% to KC162911) in *A. americanum*, and these unique sequences were deposited to GenBank (MT259333-MT259335) (Table 3). Additional testing for other tick-borne pathogens from the 2019 cohort of ticks were all negative. In 2020, we detected the same *Hepatozoon* sp. (n=12; 100% to MT259335) previous detected in 2019, *Borrelia lonestari* (n=1; 100% to AF273670), *A. phagocytophilum* (n=5; 99.4-100% to CP006617), *Ehrlichia ewingii* (n=2; 100% to U96436), and *Ehrlichia chaffeensis* (n=1; 100% to NR074500) in *A. americanum.* Sequence analysis of the 16S rRNA gene region for one *A. phagocytophilum*-positive tick (100% to MK341075) was consistent with the Ap-1 strain. For *I. scapularis*, we detected *Babesia odocoilei* (n=2; 99.76% to MH899097) and *Borrelia burgdorferi* sensu lato (n=1; 100% to AF264899). Numerous rickettsial endosymbionts (*Rickettsia amblyommatis, Rickettsia montanensis, Rickettsia* sp. TR-39, ‘*Candidatus* Midichloria mitochondrii’) were also detected from various tick species collected (Table 3).

## Discussion

In this study, we found that the *H. longicornis* populations in Virginia had similar phenology as has been previously reported in New York (Piedmonte et al., 2020; Tufts et al., 2019). To date, few studies have investigated habitat preferences for *H. longicornis* in the United States, though current data suggests that it is a habitat generalist. We detected *H. longicornis* from all habitat types during every sampling period of this two-year study in Virginia. We found *H. longicornis* on several wildlife host species, including coyote, eastern cottontail, raccoon, Virginia opossum, white-tailed deer, woodchuck, and a *Peromyscus* sp. (Table 2). In addition, we found more infections of host-seeking *H. longicornis* with the exotic pathogen *T. orientalis* Ikeda strain, as well as new reports of native pathogens (i.e., *R. felis* and *A. phagocytophilum*). These combined findings suggest that this tick may play an important and currently unrecognized role in the disease dynamics of native tick-borne pathogens warranting continued molecular surveillance to help to predict the health risks posed by this introduced species.

Our trapping efforts were focused on rodent species and meso-mammals since previous reports suggest that raccoons and Virginia opossums might be important hosts for *H. longicornis* (Beard et al., 2018; Tufts et al., 2020a; White et al., 2020). One study in New York failed to find any infested *Peromyscus* spp.; however, a later study conducted by the same group did find a single squirrel (*Sciurus carolinenesis*) to be infested with *H. longicornis* (Tufts et al., 2019, 2020a). Rodents are also important hosts for ticks and are reservoirs for numerous human tick-borne pathogens necessitating the continued surveillance of their role with *H. longicornis*. Domestic cattle are also commonly important hosts for this tick, and they do occur on the property but were not sampled because they are regularly treated with an acaracide spray. Our detections of *H. longicornis* on eastern cottontail rabbits, raccoons, woodchucks, and Virginia opossums, support results previously reported from this area and New York that this tick can use a wide range of wildlife hosts (Tufts et al., 2020a, 2019; White et al., 2020). The sampling of coyotes and white-tailed deer was opportunistic, but previous detections of *H. longicornis* on these species have also been documented (Tufts et al., 2019; USDA-APHIS-VS, 2021; White et al., 2020). Here we report an important finding of a single *H. longicornis* larva on a *Peromyscus* sp. (Table 2). This finding was surprising given experimental data showing this tick species has an aversion to smaller rodent fur (Breuner et al., 2019; Ronai et al., 2020). Importantly, this single infested animal was out of 112 *Peromyscus* sp. sampled at our site. Tufts et al. (2019) failed to find any *H. longicornis* on 190 captured *Peromyscus* sp. sampled in New York, so our detection may have been an aberrant occurrence. While some ticks depend on rodents for their earlier life stages, this does not seem the case for *H. longicornis*, as heavy infestations of larvae have been reported on Virginia opossums and raccoons (Tufts et al., 2020a; White et al., 2020).

Our seasonal density data are similar to reports from the northeastern United States, where nymphal *H. longicornis* are most active in the spring, followed by a peak in adult activity in the summer and larval activity in the fall (Figure 3) (Bickerton et al., 2020; Piedmonte et al., 2020; Tufts et al., 2019). Interestingly, we never observed a sampling period when the nymphal life stage was inactive (Figure 3). While there were gaps in our sampling periods, this suggests that in the more southern regions in the United States, there is overlap in the activity between the life stages of *H. longicornis.* This observation has been previously reported for *H. longicornis* in New Zealand and is likely due to climatic and other habitat variables within the southern United States being more favorable to multiple *H. longicornis* life stages (Heath, 2016). Since we did not detect any significant effect of microclimate variables (i.e., average wind speed, temperature, and relative humidity), more rigorous sampling of microclimate data and phenology across the recognized range of *H. longicornis* is needed to further understand the natural history of this tick and to better predict the seasonal abundance of different life stages across its range as well as potential distribution. In addition, the overlap of different *H. longicornis* life stages could complicate potential control strategies that target specific life stages. A recent study has found that environmental treatments with a pyrethroid acaricide toward the end of peak adult *H. longicornis* activity is potent enough to curb populations in the subsequent larval and nymphal life stages (Bickerton et al., 2020). Fortunately, our data suggest that the dip in nymphal activity is around this same time frame, potentially increasing this management practice’s efficacy in the southern United States.

We did not detect the exotic *Theileria orientalis* Ikeda in any screened native tick species. However, we continued to detect the pathogen in host-seeking *H. longicornis* during the 2020 sampling period, further supporting our previous results from 2019 (Thompson et al., 2020b). In addition, recent experimental work has shown that *H. longicornis* is a competent vector for this pathogen in the United States, warranting continued molecular surveillance for *T. orientalis* in *H. longicornis* in other states, especially in regions near cattle operations (Dinkel et al., n.d.). The other apicomplexan detected in *H. longicornis* was a *Hepatozoon* sp. that has previously been detected in *A. americanum* ticks from this same site and Texas (Shock et al., 2014). The vertebrate host for this parasite is currently unknown.

Two bacterial pathogens were detected in *H. longicornis.* A single tick was positive for *R. felis*, the causative agent of cat-flea typhus in humans. This pathogen also infects numerous other mammalian hosts and can be transmitted by many hematophagous arthropods, including *H. longicornis*, in low prevalence from China (Liu et al., n.d.; Pérez-Osorio et al., 2008). Because this pathogen has a broad geographic range, it is not known if it is a native or exotic strain and the genetic target is highly conserved. We detected the AP-1 strain of *A. phagocytophilum* in two *H. longicornis*. We had additional detections of *A. phagocytophilum;* however, the 16S rRNA target needed to distinguish the Ap-ha strain from AP-1 was negative for other samples positive with the *MSP-2* screening PCR. Additional studies are warranted on the role of *H. longicornis* as a vector for different *A. phagocytophilum* strains in the United States, especially since it is associated with *A. phagocytophilum* in other regions of its established range (Kim et al., 2003; Qin et al., 2018).

Notably, we did not detect any *Borrelia* or *Ehrlichia* spp. in *H. longicornis* despite detecting these pathogens in native tick species collected from the same site. The lack of *Borrelia* sp. is expected due to previous experimental work showing that *H. longicornis* is not a suitable vector for *B. burgdorferi* and other studies that have shown that rodents, the primary reservoir, are not preferred hosts (Breuner et al., 2019; Ronai et al., 2020; Tufts et al., 2019). However, a recent study from Pennsylvania did detect a single *H. longicornis* with *B. burgdorferi* s.s. using real-time PCR so additional surveillance is warranted (Price et al., 2021). Although we did not detect *Ehrlichia* spp. in *H. longicornis*, we believe that more research into their role as a vector is needed. White-tailed deer appear to be a preferred host for the tick and are important reservoirs for *E. chaffeensis* and *E. ewingii* (Lockhart et al., 1997; Tufts et al., 2019; USDA-APHIS-VS, 2021; Yabsley et al., 2002). In addition, related *Ehrlichia* spp. have also been detected in *H. longicornis* from endemic areas (Lee et al., 2005; Luo et al., 2016).

Our results show some variation in seasonal abundance of different life stages in the more southern region of this tick’s recognized distribution in the United States. In addition, a new potential host for *H. longicornis*, a *Peromyscus* sp., was documented in this study, and further investigations are needed to determine if this was an aberrant finding or if *H. longicornis* will feed on small rodents under certain circumstances. Finally, our molecular surveillance for pathogens infecting host-seeking *H. longicornis* reveals greater diversity of pathogens than previously recognized (Tufts et al., 2020b). While the role of the *Hepatozoon* sp. as a pathogen is unknown, the detections of *R. felis* and *A. phagocytophilum* AP-1 suggests that *H. longicornis* may be a vector of native pathogens circulating in our host populations. Ultimately, more investigations throughout the current range are needed to understand the ecology of *H. longicornis* and associated pathogens in the United States.

## Acknowledgments

ATT was supported by the National Science Foundation under Grant No. DGE-1545433 (UGA’s Interdisciplinary Disease Ecology Across Scales program) and through the United States Department of Agriculture Animal Plant Health Inspection Service’s National Bio- and Agro-defense Facility Scientist Training Program. The content is solely the responsibility of the authors and does not necessarily represent the official views of NSF or USDA. Partial funding was provided through Cooperative Agreements AP19VSCEAH00C004 and AP20VSCEAH00C041, Veterinary Services, Animal and Plant Health Inspection Service, US Department of Agriculture. Additional support was provided by the member states wildlife management agencies of the Southeastern Cooperative Wildlife Disease Study through the Federal Aid to Wildlife Restoration Act (50 Stat. 917) and the Ecosystems Mission Area, US Geological Survey, US Department of Interior. The authors would like to thank the landowner for granting access to his property for sampling, the many curious cow friends made over the two years, and Dr. Christopher Cleveland for his critical review of the manuscript.

